# Cardiac MRI in common marmosets revealing age-dependency of cardiac function

**DOI:** 10.1101/2020.05.18.102020

**Authors:** Amir Moussavi, Matthias Mietsch, Charis Drummer, Rüdiger Behr, Judith Mylius, Susann Boretius

**Affiliations:** Functional Imaging Laboratory, German Primate Center, Leibniz Institute for Primate Research, Göttingen, Germany; DZHK (German Center for Cardiovascular Research), partner site Göttingen, Germany; Unit of Infection Models, German Primate Center, Leibniz Institute for Primate Research, Göttingen, Germany; Department of Laboratory Animal Science, German Primate Center, Leibniz Institute for Primate Research, Göttingen, Germany; Platform Degenerative Diseases, German Primate Center, Leibniz Institute for Primate Research, Göttingen, Germany

## Abstract

The aim of this study was to establish a feasible and robust magnetic resonance imaging protocol for the quantitative assessment of cardiac function in marmosets and to present normal values of cardiac function across different ages from young adult, middle-aged, to very old clinically healthy animals.

Cardiac MRI of 33 anesthetized marmosets at the age of 2-15 years was performed at 9.4 T using IntraGate-FLASH that operates without any ECG-triggering and breath holding. Normalized to post-mortem heart weight, the left ventricular end-diastolic volume (LV-EDV) was significantly reduced in older marmosets. The LV end-systolic volume (LV-ESV) and the LV stroke volume (LV-SV) showed a similar trend while the LV ejection fraction (LV-EF) and wall thickening remained unchanged. Similar observations were made for the right ventricle. Moreover, the total ventricular myocardial volume was lower in older monkeys while no significant difference in heart weight was found.

In conclusion, IntraGate-FLASH allowed for quantification of left ventricular cardiac function but seems to underestimate the volumes of the right ventricle. Although less strong and without significant sex differences, the observed age related changes were similar to previously reported findings in humans supporting marmosets as a model system for age related cardiovascular human diseases.

## Introduction

In the recent years a small New World non-human primate (NHP), the common marmoset (*Callithrix jacchus*), has become increasingly popular in preclinical biomedical research, not least because of the recent advantages in generating transgenic marmoset offspring [1, 2]. Marmosets have been extensively used as a model system for studying reproduction, infectious diseases, brain functions and aging [3–6]. Moreover, obesity and diabetes, two common risk factors for cardiovascular diseases in humans, have been intensively explored in marmosets [7–10]. Using radionuclide angiography, Charnock et al. [10, 11] showed that nutrition high in dietary fat can significantly influence the composition of cardiac membranes towards increased level of fatty acids. In addition, those marmosets showed an increased heart rate together with an increased left ventricular ejection fraction.

Magnetic resonance imaging (MRI) is currently one of the main imaging techniques for the assessment of cardiac function and morphology in human. Recently, cardiovascular MRI has also been successfully used in macaques [12–14]. For instance, cardiac effects of spontaneous type 2 diabetes [13] and the effects of myocardial infarction [14] on the cardiac performance have been studied in adult cynomolgus and rhesus macaques. At present, however, there is a lack of MR imaging protocols and reference values of cardiovascular MRI in marmosets.

In our rapidly aging societies, suitable animal models most closely resembling human aging are urgently required. Besides similarity to humans and other NHPs [15, 16], particular advantages of the marmoset model in aging research is a relatively short generation time, small size and availability of genetically modified models. Several groups have shown age-related changes in the pathology of common marmosets but have focused mainly on the central nervous system [5–7]. Two post-mortem studies looking for the cause of death in two large colonies showed a significant higher prevalence of cardiac fibrosis in aged marmosets [5, 17], but only little is known about the consequences of increased age on cardiac function in this species.

The aim of this work was to establish a robust and feasible MRI protocol to measure the cardiac function of marmosets and to present average values of cardiac function across clinically healthy animals of different ages ranging from young adults (2 years) up to very old marmosets (15 years). Difficulties of, and possible solutions for, carrying out cardiovascular MRI of marmosets will be discussed.

## Results

### Image Quality

Short-axis images were acquired during free breathing using a navigator based IntraGate-FLASH. The quality of the data sets was investigated by visual inspection and blinded rated using a 5 point-score (1 for poor and 5 for very good quality). Based on that 4 out of 33 animals were excluded from further data analysis due to a score lower 3 (2x score 1, 1x score 1.5, 1x score 2.5). In addition, one data set had to be excluded due to a circumscribed signal loss in region of the right ventricle. The quality score of all included animals was on average 4.5 ± 0.6 (range: 3-5). Figure 1a shows the respective image of a mid-ventricular slice (diastolic phase) of the excluded animals compared to an animal with the highest quality score (Animal A). Respiratory artifacts (Animal B), no visible ventricular contraction (Animal C and D), low blood-myocardium contrast (Animal D and E) and fuzziness (Animal D) were the main artifacts that significantly reduced the diagnostic value of the images.

**Figure 1:**
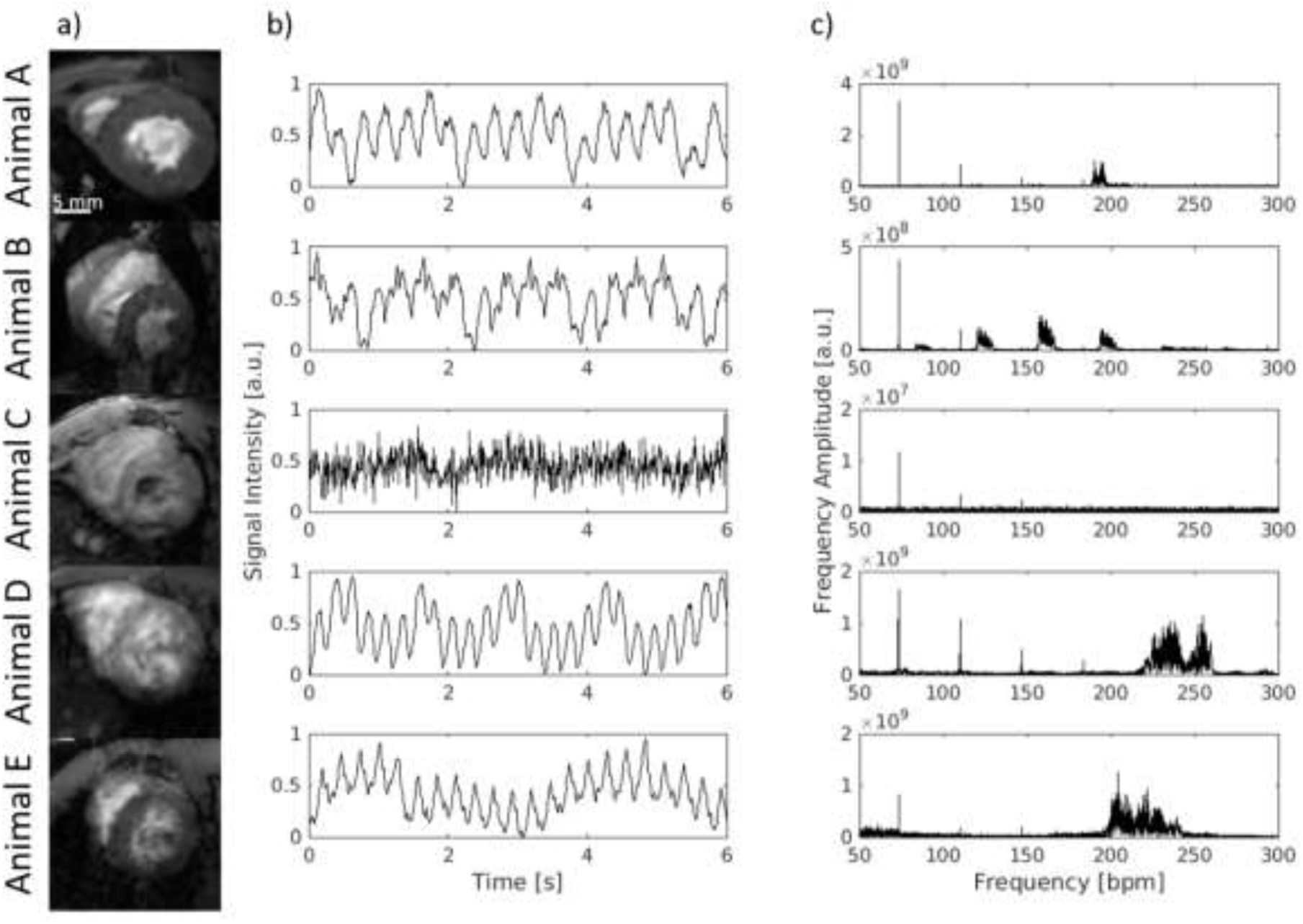
Achieved image quality and navigator signal. a) Only images with at least good quality (e.g. animal A, score = 5) were selected for further quantification whereas images with low quality (scoring < 3) were excluded from further quantification (animal B-E). Respiratory artifacts (B), no visible ventricular contraction (B, C), low blood-myocardium contrast (C, E) and fuzziness (D) are the main image artifacts which significantly reduced the diagnostic value of the images. All images are in the diastolic phase. b) Time course of the navigator signal of the excluded animals (B-E) compared to a high-score animal (A): Signal changes due to cardiovascular movement are visible although to different extent. c) Fourier analysis of the navigator signal: the sharp peak at 35 bpm represents the respiration whereas the widespread peak accumulation around 200-250 bpm corresponds to the cardiac rate.

Parts of these artefacts had their origin in a corrupted navigator signal as illustrated in Figure 1b. Essential for the reconstruction of images with superior quality was the appearance of a relatively sharp and compact frequency band in the range of the cardiac rate as shown by the Fourier transformation of the navigator signal of a high-score-image (Fig. 1c; Animal A).

The cardiac rates observed under anesthesia were in the range of 150 and 250 bpm. Fluctuations of this rate throughout data acquisition led to widespread peaks in the frequency domain (Animal D and E) and as a result to a clearly reduced image quality. Similarly, additional signal fluctuations (Animal B) led to an insufficient image quality. A lack of rhythmic signal fluctuations (Animal C) resulted in no visible ventricular contraction on the reconstructed dynamic images. Consequently, these corruptions of the navigator signal led to an exclusion of the data sets and may be used in future for automatic quality control.

Changes of the signal intensity related to respiratory movement could be detected on all navigator signal curves. Due to the respiratory frequency given by the artificial ventilation frequency analyses revealed always a sharp peak around 35 bpm. In addition, in most of the cases harmonics of the respiration were visible at multiples of 35 bpm.

Figure 2 exemplarily shows diastolic and systolic images of two animals (Animals F and G) with good image quality throughout all acquired slices. In particular, papillary muscles and trabecularization were well resolved. Although residual flow artifacts and fuzziness may still be visible on these images, the achieved quality was sufficient for quantitative analysis of cardiac function.

**Figure 2:**
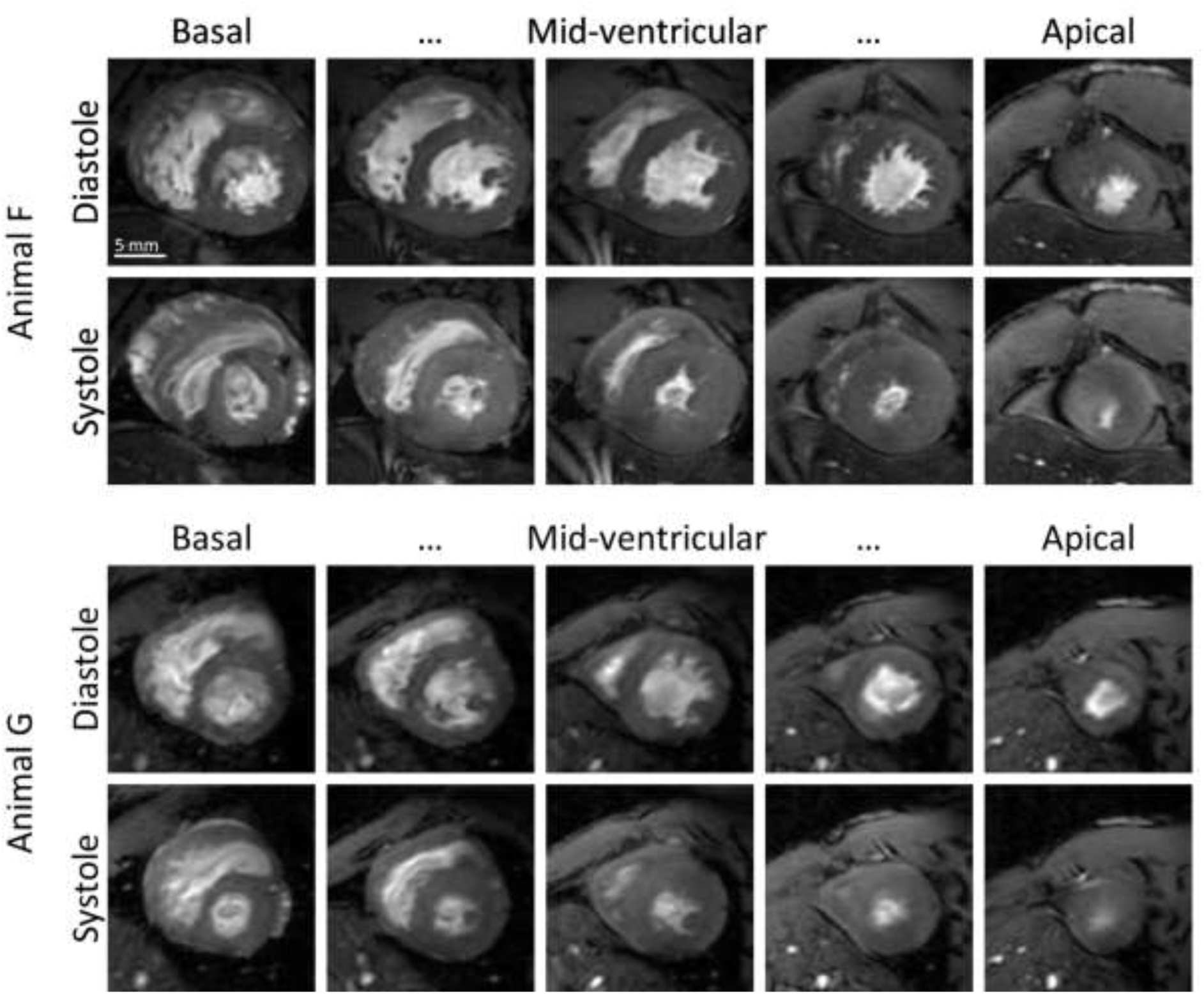
Diastole and Systole. Diastolic and systolic images of the heart of two marmosets (Animal F and G) with the highest quality score of 5. In particular, papillary muscles and trabecularization were well resolved.

### Cardiac Mass and Function

All marmosets were grouped by age into three groups for further analyses (young adult (< 5 years), middle-aged (5 – 9 years), and senescent (>9 years). The respective group compositions as well as mean values of heart rate, body weight, and post-mortem heart weight are shown in Table 1. Marmosets, in contrast to most other primates, exhibit no somatic sex-dimorphism. Moreover, a two-way ANOVA (with age group and sex as independent variable) revealed no significant differences between the sexes in none of the analyzed parameters. For these reasons cardiac parameters obtained from male and female marmosets where pooled together (Table 2).

**Tab. 1:** Marmosets included in the study: Age groups, mean values of heart rate, body weight, and heart weight.

| Parameter | Age groups |  |  | Female | Male | All | p-value<br>(one-way<br>ANOVA,<br>age) |
| --- | --- | --- | --- | --- | --- | --- | --- |
|  | 23-58<br>Months | 63-106<br>Months | 110-168<br>Months |  |  |  |  |
| Animals (n)<br>Female/Male | 10<br>5/5 | 9<br>6/3 | 10<br>3/7 | 14 | 15 | 29 | - |
| Age (months) | 36 ± 14 | 91 ± 14 | 138 ± 18 | 82 ± 42 | 94 ± 47 | 88 ± 45 | - |
| HR (bpm) | 188 ± 19 | 193 ± 29 | 203 ± 40 | 194 ± 30 | 194 ± 32 | 195 ± 31 | 0.51 |
| Weight (g) | 421 ± 54 | 402 ± 39 | 395 ± 43 | 419 ± 37 | 398 ± 53 | 406 ± 45 | 0.29 |
| Heart Weight (g) | 2.46 ± 0.37 | 2.63 ± 0.32 | 2.64 ± 0.38 | 2.61 ± 0.35 | 2.55 ± 0.37 | 2.58 ± 0.36 | 0.20 |

All data given as arithmetic mean and standard deviation. Significance level $p < 0.05$ .

**Tab. 2:** Left and right ventricular Function.

| Parameter | Age groups |  |  | Female | Male (all) | All | p-value<br>(one-way ANOVA,<br>age) |
| --- | --- | --- | --- | --- | --- | --- | --- |
|  | 23-58<br>Months | 63-106<br>Months | 110-168<br>Months |  |  |  |  |
| TV-MV (μl) | 1506 ± 247 | 1469 ± 187 | 1353 ± 166 | 1417 ± 170 | 1456 ± 235 | 1437 ± 203 | 0.22 |
| LV-MV (μl) | 831 ± 107 | 908 ± 135 | 819 ± 105 | 844 ± 128 | 866 ± 112 | 851 ± 119 | 0.26 |
| LV-EDV (μl) | 817 ± 129 | 731 ± 181 | 720 ± 104 | 793 ± 183 | 740 ± 113 | 755 ± 142 | 0.14 |
| LV-ESV (μl) | 352 ± 47 | 326 ± 93 | 308 ± 52 | 352 ± 96 | 319 ± 48 | 328 ± 67 | 0.18 |
| LV-SV (μl) | 465 ± 109 | 405 ± 1116 | 411 ± 99 | 441 ± 120 | 421 ± 98 | 427 ± 107 | 0.32 |
| LV-EF (%) | 56.3 ± 6.8 | 55.2 ± 7.0 | 56.8 ± 7.0 | 55.3 ± 7.4 | 56.5 ± 6.2 | 56.1 ± 6.8 | 0.87 |
| WallTh (mm) | 1.07 ± 0.37 | 1.06 ± 0.40 | 1.00 ± 0.22 | 1.02 ± 0.38 | 1.06 ± 0.28 | 1.04 ± 0.32 | 0.88 |
| FracWallTh(%) | 59.4 ± 22.2 | 51.1 ± 23.5 | 53.0 ± 13.7 | 54.4 ± 21.8 | 54.8 ± 18.4 | 54.6 ± 19.8 | 0.65 |
| RV-MV (μl) | 672 ± 144 | 561 ± 68 | 534 ± 108 | 581 ± 86 | 593 ± 154 | 587 ± 123 | 0.03* |
| RV-EDV (μL) | 620 ± 139 | 490 ± 87 | 452 ± 118 | 535 ± 152 | 502 ± 116 | 519 ± 134 | 0.01* |
| RV-ESV (μL) | 355 ± 70 | 292 ± 68 | 257 ± 63 | 299 ± 86 | 301 ± 69 | 300 ± 77 | 0.01* |
| RV-SV (μL) | 265 ± 86 | 198 ± 45 | 195 ± 86 | 236 ± 81 | 201 ± 76 | 219 ± 79 | 0.10 |
| RV-EF (%) | 42.0 ± 7.5 | 40.7 ± 8.5 | 41.6 ± 12.4 | 43.8 ± 8.7 | 39.1 ± 9.9 | 41.5 ± 9.4 | 0.96 |

All data given as arithmetic mean and standard deviation. Significance level $p < 0.05$ .

#### Cardiac Mass

The weight of the hearts, measured post-mortem and including ventricles and atria, ranged from 1.85 g to 3.13 g (2.6 ± 0.4 g, mean ± standard deviation). This weight was positively correlated with the body weight of the animal (Pearson’s r = 0.54, p < 0.01) and the ventricle myocardial volume measured by MRI (LV-MV: Pearson’s r = 0.61, p < 0.01 and TV-MV: Pearson’s r = 0.48, p < 0.01; Fig. 3a-c). With higher age the heart weight and the heart rate showed a tendency to higher values while the body weight seemed to slightly decrease (Fig. 3d-f). But none of these observations reached statistical significance (p (body weight) = 0.29, p (heart rate) = 0.51, p (post mortem heart weight) = 0.51).

**Figure 3:**
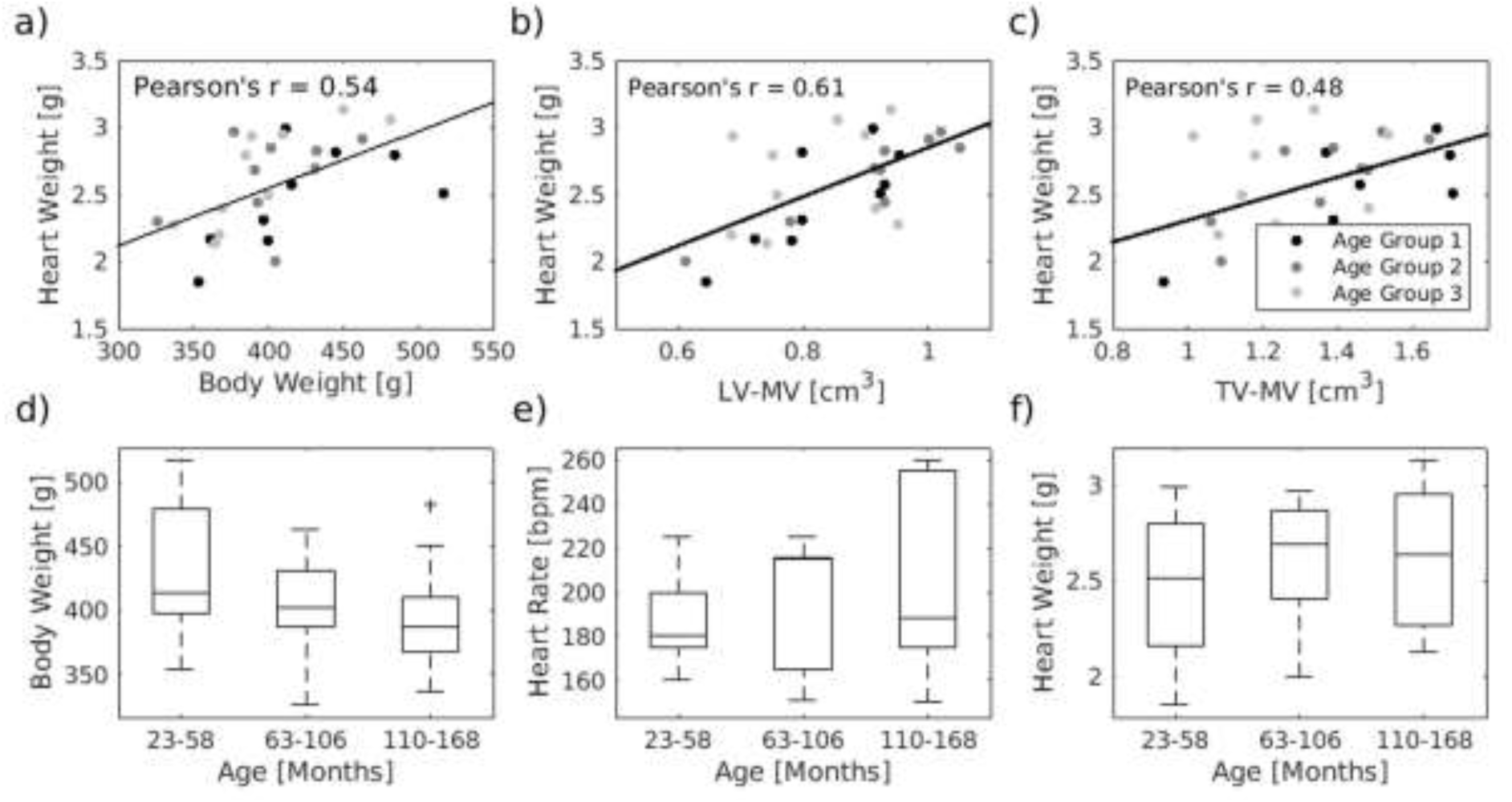
Age dependency of physiological parameter. (a-c) The weight of the hearts correlates with the body weight of the animals, the left ventricular myocardial volume and the total muscle volume (p < 0.01 for all). Body weight (d), heat rate (e) and post-mortem weight of the heart (f) of the marmosets were almost equally distributed between the 3 age groups (*n* = 10/9/10). The central line of the box plot indicates the median, the edges are the 25^th^ and 75^th^ percentile and the outliers are visualized by the ‘+’ symbol.

Significant age related effects could, however, be observed for the ventricle myocardial volume. A reduction with age involved mainly the right ventricle (p = 0.03) and became particular evident when normalizing the myocardial volume by the respective post-mortem heart weight (Fig. 4a, c). This increase in myocardial density (Fig. 4d) was already significant when comparing the first and second age group (t-test, p < 0.003).

**Figure 4:**
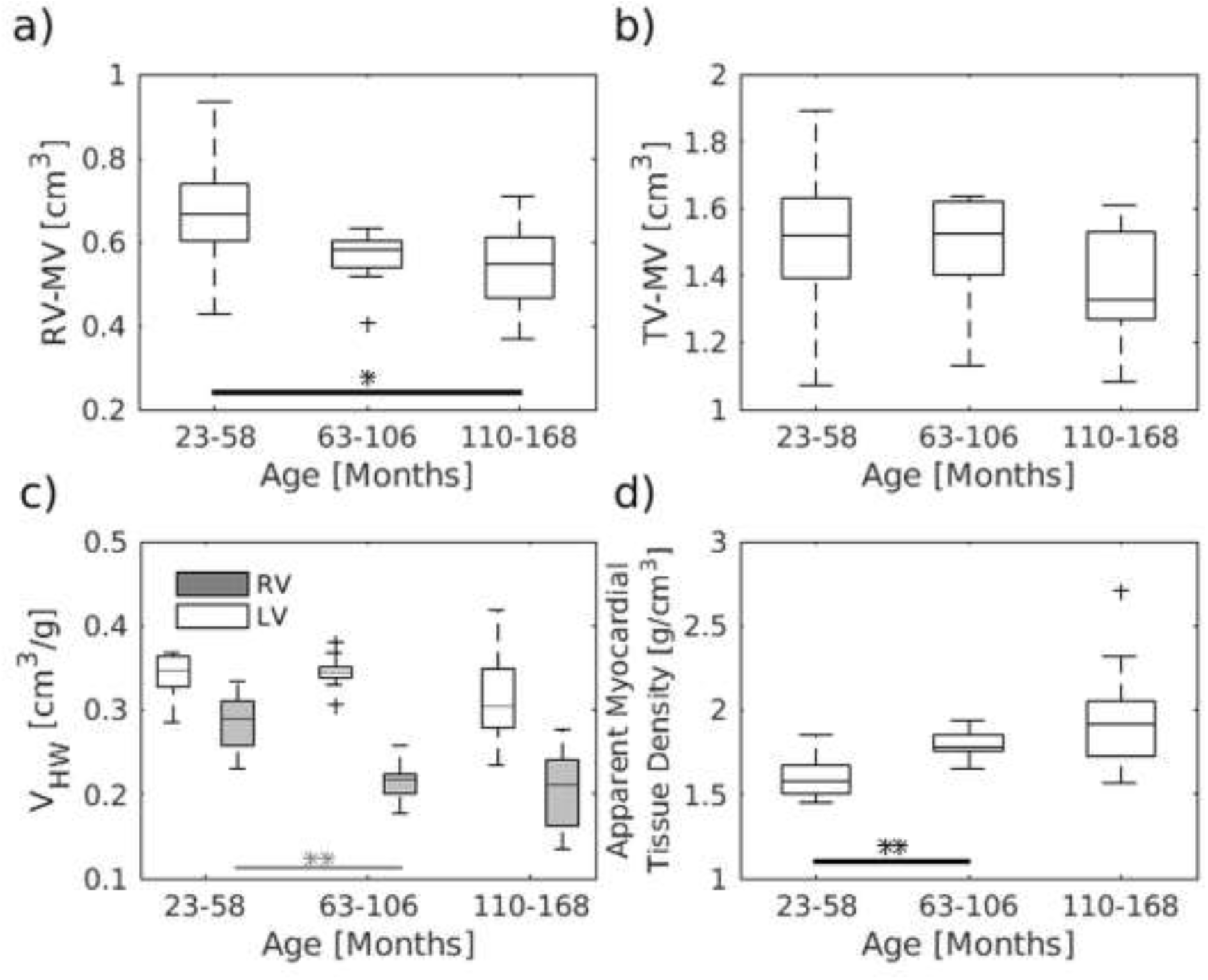
Age dependency of ventricle myocardial volume. The right and total volume of the ventricle muscle were decreased with age (a, b). This became even more evident when normalizing the volumes to the post-mortem weight of the heart (c) and involved in particular the right ventricle (a, c). This volume lost was accompanied with an increase in the apparent myocardial tissue density of the myocardium (d). The data were compared using one-way ANOVA, followed by a t-test (*P < 0.05, **P < 0.01, ***P < 0.001). The central line of the box plot indicates the median, the edges are the 25^th^ and 75^th^ percentile and the outliers are visualized by the ‘+’ symbol. *n* = 10/9/10 for all graphs.

#### Function of the left ventricle

The stroke volume (SV) of the left ventricle averaged over all analyzed marmosets was 427 ± 107 μl in total and corresponded to 1.05 ± 0.23 μl/g body weight. The mean respective end-diastolic volume (EDV) and end-systolic volume (ESV) were 755 ± 142 μL and 328 ± 67 μL resulting in an average ejection fraction (EF) of 56 ± 7 % (range: 41.5 - 69.0 %). Using absolute values one-way ANOVA revealed no significant age effect for none of the acquired parameters of left ventricle function (Left column of Fig. 5 and Tab. 2).

**Figure 5:**
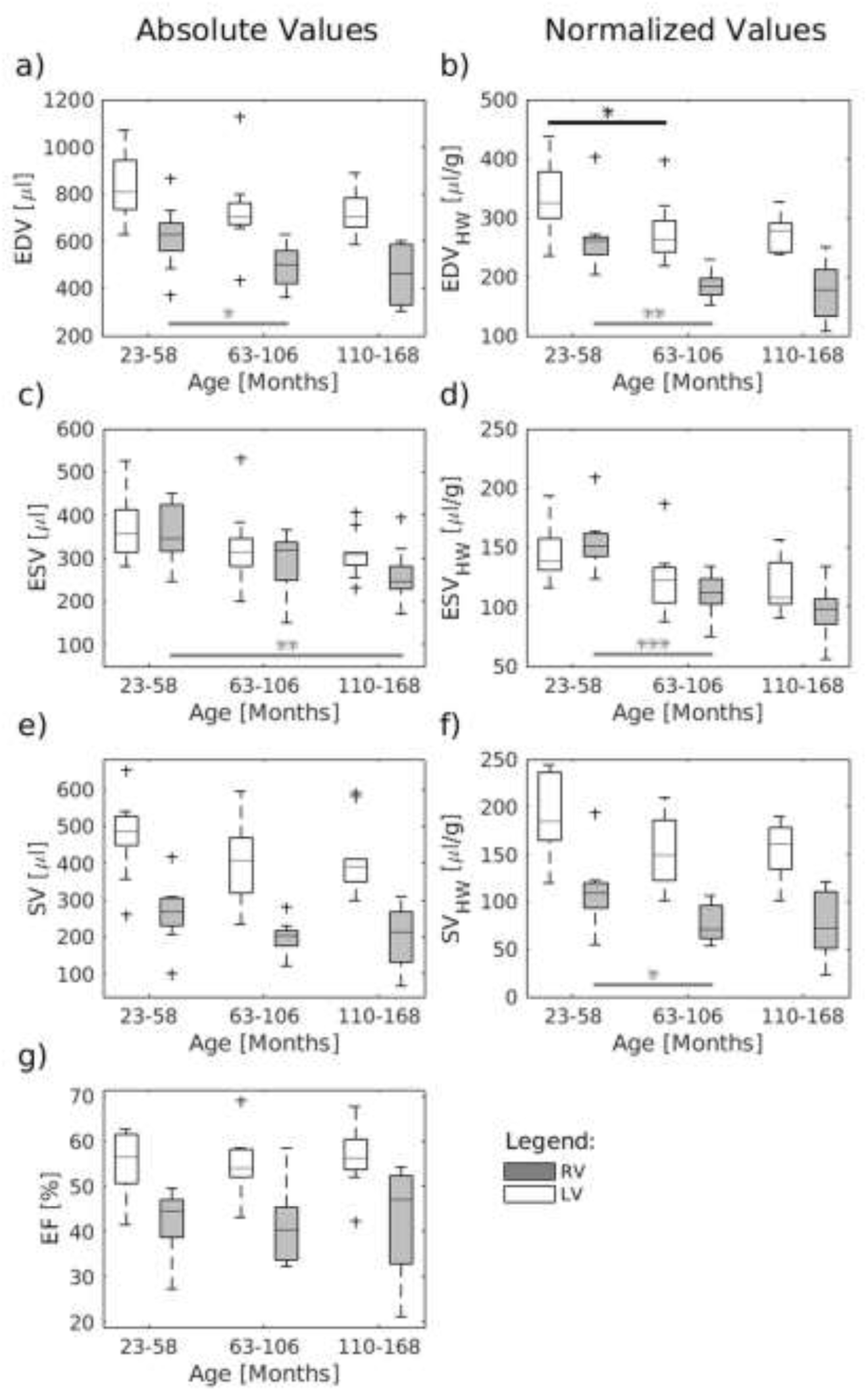
Age dependency of cardiac function. EDV, ESV and SV showed for both the left and right ventricle a trend to lower values in older marmosets. However, only RV-EDV and RV-ESV revealed a significant age effect. When correcting for the weight of the heart the LV-EDV showed a significant reduction with age. The SV relative to the heart weight also revealed a tendency to lower values in old marmosets but missed the level of significance for both left and right ventricle. The EF was almost equal within the age group. The data were compared using one-way ANOVA, followed by a t-test (*P < 0.05, **P < 0.01, ***P < 0.001). The central line of the box plot indicates the median, the edges are the 25^th^ and 75^th^ percentile and the outliers are visualized by the ‘+’ symbol. *n* = 10/9/10 for all graphs.

This was still the case when normalizing these volumes to body weight or body surface area (BSA). However, when correcting for the weight of the heart (Table 3) the EDV showed a significant reduction with age (right column of Fig. 5, p = 0.02). For both, ESV (p = 0.07) and SV (p = 0.08) a tendency to lower values could be observed but missed the level of significance.

**Tab 3:**
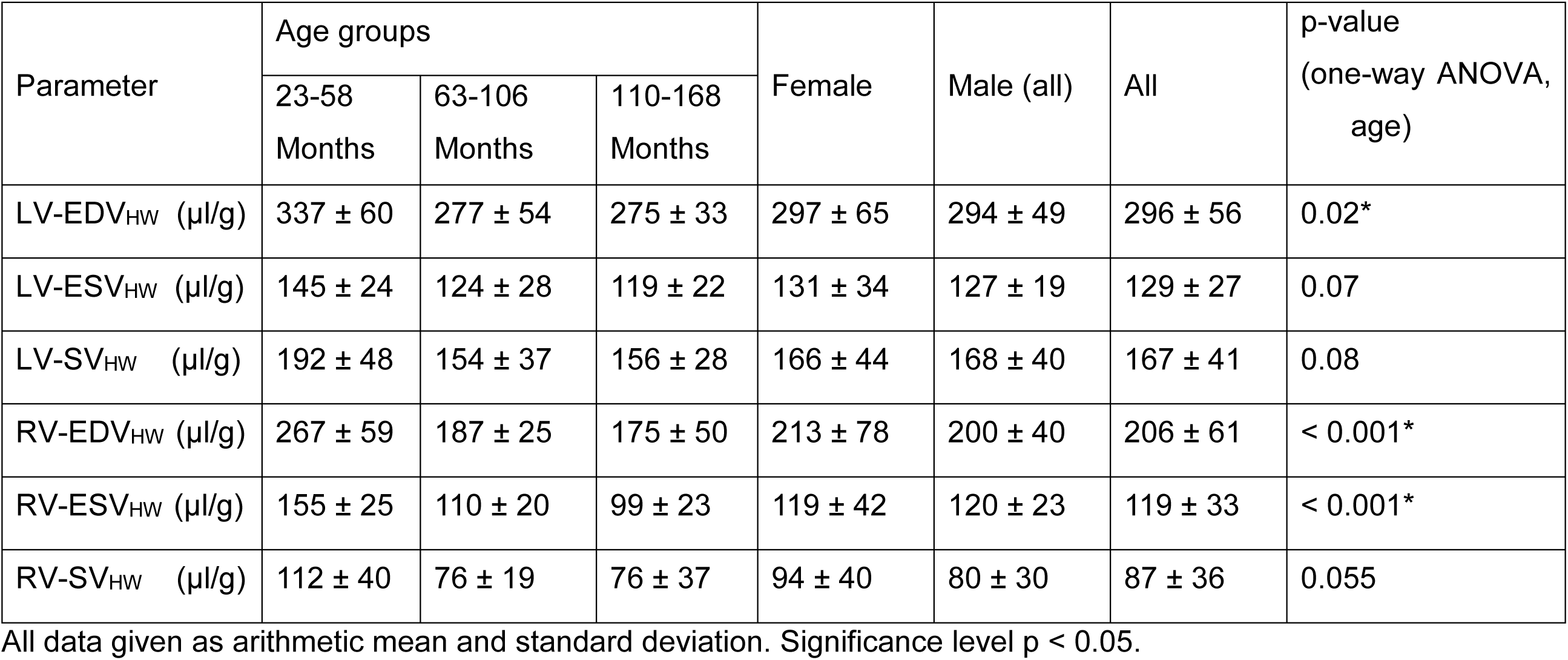
Left and right ventricular volumes relative to the post-mortem weight of the heart.

The EF was, as expected, positively correlated with the absolute and fractional wall thickening (Supplementary Figure 1, Pearson’s r = 0.76 and 0.75, respectively). The thickening of the wall showed a significant region dependency (p < 0.002) exhibiting the highest values in the inferolateral and anterolateral segments and the lowest values in the anterior and anteroseptal segment, independent of age, and weight of the animal (Supplementary Table).

#### Function of the right ventricle

The EDV and ESV of the right ventricle averaged over all marmosets were 519 ± 134 µl and 300 ± 77 µl, respectively. The resulting mean SV of the right ventricle equaled with 219 ± 79 µl just less than half of the left ventricle SV. This indicates systematic errors in determining the absolute right ventricle volumes since it is usually more difficult to determine than those of the left ventricle. Assuming, however, that these methodical errors apply almost equally to all data sets, EDV and ESV showed a significant age effect exhibiting lower values at higher age (left column of Fig. 5, both p = 0.01). This became even more evident when correcting for heart weight (right column of Fig. 5, both p < 0.001). The SV relative to the heart weight also revealed a tendency to lower values in old marmosets but missed the level of significance (p = 0.055). As seen for the left ventricle, the EF of the right ventricle remained constant with age. None of the parameters showed significant differences between male and female monkeys.

## Discussion

To the best of our knowledge this study is the first report of cardiovascular MR in common marmosets. The left and right ventricular cardiac functionality of 29 clinically healthy appearing common marmosets were assessed and the respective values were reported related to the age and the sex of the monkeys. Both left and right ventricular function showed significant age effects although to different extent. No significant differences between male and female monkeys could be observed. The findings will be discussed in the following in more detail.

Utilizing a navigator based IntraGate Flash sequence, a method that has been used so far mainly in rodents [18, 19], cardiac MRI could be performed without the necessity of breath holding and ECG recording. The image quality was satisfactory, only 4 of the 33 acquired data sets had to be excluded from further analysis because of insufficient or too much fluctuating navigator signals. A situation where this method failed most severely was a fluctuation of the cardiac rate during the relative long acquisition time of 23 minutes. For comparison, CINE MRI in human and macaques requires about 15s per slice. Here further methodological development is certainly required (see below). However, in all 29 animals included, a very good delineation of the cardiac wall and a sufficient visualization of papillary and trabecular muscles were achieved.

The volume of the ventricular myocardium and likewise the post-mortem heart weight were significantly correlated with the body weight of the marmoset. The average LV-SV of 1.05 ± 0.2 μl/g body weight corresponds to what has been reported in macaques [12, 14] and humans [20, 21]. In contrast to humans, in whom on average the mass and size of the heart is lower in women compared to men [20], male and female marmosets showed comparable heart weights. Together with the similarity in body weight between sexes, this observation extend the finding that marmosets, in contrast to most other primates, exhibit no somatic sex-dimorphism, neither externally nor internally. Based on that, we pooled the data of both sexes together. More detailed analysis of subtle sex differences may require a larger number of marmosets per group.

Aging is a main risk factor for cardiovascular diseases. A variety of studies investigated the effect of age on cardiac anatomy and function in humans. The Multi-Ethnic Study of Atherosclerosis (MESA), for instance, that investigated men and women between the age 45 and 84 and free of cardiovascular diseases, revealed progressively reduced EDV and SV in older age whereas EF remained almost unchanged [20]. In addition, Maceira and colleges [22] reported a reduction of the LV-ESV in elderly people. Between the age of about 20 to 80 years volume reductions were likewise found for the right ventricle including EDV, ESV, and SV [23].

Taking an average life expectancy of about 12 years for marmosets in captivity, the age range of the marmosets included in this study corresponds to about 18 to 80 years in humans. Even though previous studies have reported higher risk of myocardial fibrosis [5, 17] and increased blood pressure [24, 25] with increasing age in marmoset, we observed only relatively mild disturbances of the cardiac function with age.

In particular, the absolute values of the left ventricular volume showed no significant age effect but a relatively large variation between the animals. To compensate for physiological variations in human these values are commonly reported normalized to BSA or metabolic body weight. None of these normalizations changed the result in our marmoset study significantly. Interestingly, however, when correcting for post-mortem heart weight, all left ventricle volumes showed a tendency of reduction with age, with EDV reaching the level of significance. In addition, the EDV and ESV of the right ventricle were reduced with age, reaching the level of significance already for the absolute values. Likewise previously reported for humans [20], this reduction in ventricular volumes did not lead to a reduction of EF or wall thickening. Both, EF and wall thickening remained preserved in elderly marmoset, at least under rest. Another parameter that has been put into the context of aging is the left ventricular mass (LVM). It is defined as the volume of the myocardial muscle multiplied by 1.05 [26]. Follow up measurements in humans revealed an increase in LVM with age in men and a mild decrease in women. Cross sectional, however, the LVM was decreased in elderly asymptomatic humans of both sexes [21]. The latter is very much in line with our observations in marmosets. Moreover, since we also know the post-mortem heart mass of these animals, it became clear that this reduction of myocardial volume went along with an increase in tissue density by about 23% (group 1 versus group 3). Thus, the assumption of a fixed muscle density in order to calculate the LVM is misleading and it may be more advisable to use the ventricular muscle volume (LV-MV) instead.

The observed changes in myocardial density fit nicely to reports on cardiac histopathology in marmosets and humans showing that tissue remodeling and diffuse interstitial fibrosis are main characteristics in cardiac life course. Future MRI studies should certainly utilize fibrosis sensitive techniques including gadolinium enhanced MRI and mapping of T1 relaxivity, which may in particular help to unravel the time course of tissue remodeling in follow up studies.

A limitation of this study became apparent by the lower stroke volumes for the right ventricle compared to the left. Under physiological condition, these volumes should of course be similar. Differences in LV-SV and RV-SV, however, have been reported by several MRI studies [27, 28]. The reasons for that discrepancy are most likely of technical nature. In particular, the oblique orientation of the tricuspid valve compared to the mitral valve which is usually used to align the short-axis view, hampers a correct detection of the atrio-ventricular valve plane of the right ventricle. Because of this a precise determination of the right ventricle volume is a well-known challenge in cardiac MRI and the method applied in this study does not seem to master this challenge. This systematic error makes it impossible to determine the exact absolute RV volumes. Interestingly, however, the difference between left and right SV was independent of age (p = 0.97). Thus, cautiously drawing conclusions about the relative differences obtained from the marmosets within the study may still be appropriate.

Improvements for future studies should certainly include the use of multichannel-coils. Parallel imaging techniques can then be used to speed up the acquisition. The relatively long scan time was one of the main sources for the observed image artefacts. Moreover, a better spatial resolution including thinner slices will certainly help to reduce the underestimation of the RV-SV in future studies. In addition, more complex navigator techniques which allow the detection of arrhythmias may increase the success rate of the data acquisition. Further development will include the usage of non-Cartesian k-space techniques like radial or spiral trajectories to increase the spatiotemporal resolution and to reduce respiratory artifacts.

In conclusion, the observed age related changes in marmoset were rather similar to previously reported findings in clinically healthy humans although to a lower extent. These changes involved a reduction of the ventricular volumes with increase in age while cardiac efficiency largely remained unaffected. Remodeling processes of the myocardium seem already to occur in middle aged marmosets including a reduction of myocardial volume by increased tissue density. Taking into account the cardiovascular similarities that share marmosets with humans [15, 16] and the recent achievements in genetic manipulation of these monkeys [1, 2], these findings strongly support marmosets for the use as model system for age related cardiovascular diseases.

## Methods

### Animals

Experiments were conducted on 33 adult, clinically healthy common marmoset monkeys (*Callithrix jacchus*) with an age ranging from two to 15 years (7.4 ± 3.8 years) and an average weight of 408 ± 46 g. The study cohort consisted of 15 females and 14 males. The animals were grouped by age as follow: young adult (< 5 years), middle-aged (5 – 9 years), and senescent (>9 years) [5, 29].

Based on the achieved image quality data sets of 3 animals had to be excluded from further analyses resulting in an age distribution within the group of 1.9 – 4.8 years (young adult), 5.3 – 8.8 years (middle-aged), and 9.1 – 14 years (senescent).

All animals were raised and kept in accordance with the German Animal Welfare Act. All experiments were carried out with the approval of the ethics committee of the Lower Saxony State Office for Consumer Protection and Food Safety (33.19-42502-04-17-2496) and in accordance to the guidelines from Directive 2010/63/EU of the European Parliament on the protection of animals used for scientific purposes.

Marmosets were initially anesthetized with a mixture of alfaxalone (12 mg/kg, Alfaxan, Jurox) and diazepam (3 mg/Kg, Ratiopharm). This was followed by 0.05 ml glycopyrronium bromide (0.2 mg/ml, Robinul, Riemser, Biosyn) per animal to prevent secretion, maropitant (1 mg/kg, Cerenia, Pfizer) as an antiemetic, and meloxicam (0.2 mg/kg, Metacam, Boehringer Ingelheim) as anti-inflammatory analgesic. Anesthesia was maintained with 0.9 ± 0.3 % isoflurane in a mixture of oxygen and ambient air as required. Marmosets were mechanically ventilated via an endotracheal tube for the duration of the measurement (respiration rate = 35 bpm). The body temperature was kept constant by blankets filled with warm water. Physiological parameters, such as rectal temperature, heart rate, chest movement and peripheral arterial oxygen saturation were continuously monitored during experimentation (ERT Control/Gating Module 1030, SA instruments).

In order to perform MRI anesthetized marmosets were positioned in the sphinx position in a custom-made MRI-compatible stereotaxic frame to minimize movement artifacts. The ear bars served additionally as hearing protection and were dabbed with a lidocain containing ointment (Emla 5%, AstraZeneca) for local anesthesia. Application of eye ointment (Bepanthen AS, Bayer) prevented the eyes from drying out during anesthesia.

At the end of the study animals were deeply anaesthetized with a combination of ketamine (i.p.,50mg/kg, Keatmin 10%, WDT), xylazine (i.p.,10mg/kg, Xylariem 2%, Ecuphar) and atropine (i.p.,1mg/kg, Atropinsulfat, Dr. Franz Koehler Chemie GmbH) followed by an intraperitoneal administration of pentobarbital (150-200mg/kg). A broad spectrum of organs including the heart was collected for further analyses.

### MRI

Cardiovascular MRI were obtained on a 9.4 Tesla small animal MRI-system (BioSpec 94/30, Bruker BioSpin MRI GmbH, Ettlingen, Germany) equipped with a 330 mT/m gradient system (BGA-20S, Bruker BioSpin MRI GmbH, Ettlingen, Germany). A custom made ellipsoid single-loop receive coil with a diameter of 38 mm x 35 mm (O-HLE-94, Rapid Biomedical, Rimpar, Germany) was positioned underneath the chest for optimal coverage of the heart.

For slice positioning, low-resolution (1.55 × 1.55 × 8 mm^3^) ECG-gated T1-weighted images were obtained during free breathing covering the heart in three orientations (2-chamber, 4-chamber and short-axis). To quantify the cardiac function, short-axis images were acquired during free breathing using a navigator based IntraGate-FLASH sequence with following parameters: 40 navigator points, repetition time (TR) = 48 ms, echo time (TE) = 2 ms, flip angle = 12°, Field of View (FOV) = 55 × 55 mm^2^, acquisition bandwidth = 85 kHz, echo position = 33.33%, spatial resolution = 0.215 × 0.215 × 1 mm^3^, 2 × 2 in-plane interpolation, slice gap = 1.5 mm, 5 slices (from apex to base) and 200 repetitions resulting in a total acquisition time of approximately 23 minutes.

Cardiorespiratory rates were extracted from the navigator signal using a Fourier analysis and used as gating signal for retrospective image reconstruction [18]. To get an appropriate coverage of the cardiac cycle, 16 cardiac phases per slice were reconstructed. The achieved image quality was investigated by visual inspection and by scoring the following criteria: artifacts due to breathing, blood-myocardium contrast and detection of diastole and systole. The score ranged from 1 = poor quality to 5 = very good quality. Images scored less than 3 were excluded from further analysis.

### Assessment of cardiac function

The left and right ventricle function were evaluated using short-axis images and the freely available software package Segment (Version 2.0 R6435, Medviso, Lund, Sweden) [30]. The end-diastolic and the end-systolic phase were selected as the images with the highest and lowest blood volume of a mid-ventricular slice, respectively. Endocardial and epicardial contours were manually drawn from base to apex of the heart in both cardiac phases. The end-diastolic volume (EDV) and the end-systolic volume (ESV) were calculated using Simpson’s method of disks [31]. The stroke volume (SV) and the ejection fraction (EF) were defined as the difference between EDV and ESV and the relation of SV and EDV, respectively. The myocardial volumes (MV) of the left and right ventricle (excluding the atria) were defined as the volume difference of the corresponding epicardial and endocardial volumes of the ventricles. The total ventricular myocardial volume (TV-MV) was defined as the sum of LV-MV and RV-MV. The left ventricular diastolic/systolic wall thickening (DWallTh/SWallTh), absolute wall thickening (WallTh) and the fractional wall thickening (FracWallTh) were calculated using a six sector bullseye of the three mid-ventricular slices. The six sectors were defined as anterior, anteroseptal, inferoseptal, inferior, inferolateral, and anterolateral (Supplementary Figure 2).

To compensate for the heterogeneous size of the animals, following parameters were used to normalize the volumetric parameter: the post-mortem heart weight (HW), the body weight (BW), and the body surface area (BSA). The latter was calculated according to the equation proposed for macaques [32–34]. The apparent myocardial tissue density was defined as the ratio of post-mortem heart weight and the total ventricular myocardial volume measured by MRI.

### Statistical analysis

All values reported are given as arithmetic mean ± standard deviation. Statistical analysis was performed using Matlab (R2015a, MathWorks, Natick, USA). Unless otherwise stated one-way ANOVA with post hoc t-test was used to calculate significant age differences between the groups. A value of P < 0.05 was considered as statistically significant, because this is the first explorative study on the topic we did not adjust for multiple testing.

## Acknowledgements

We thank Hasti Ghasemipour, Salim Ansari, Kerstin Fuhrmann, and Kristin Kötz for their technical assistance in data acquisition and analyses.

## Author contributions

All authors conceived and designed the study. AM, JM and SB performed the experiments and collected the data. AM and SB performed the analyses and wrote the manuscript. All authors participated in the critical manuscript revision, read and approved the manuscript in its final version.

## Data availability

The datasets generated and analyzed during the current study are available from the corresponding author on reasonable request.

