## Supplemental File for "Cardiac MRI in common marmosets revealing age-dependency of cardiac function"

**Supplementary Figure 1: Wall thickening:** Both the absolute and fractional wall thickening correlate with the estimated LV-EF (Pearson's r = 0.76 and 0.75 with $p < 0.001$ for both).

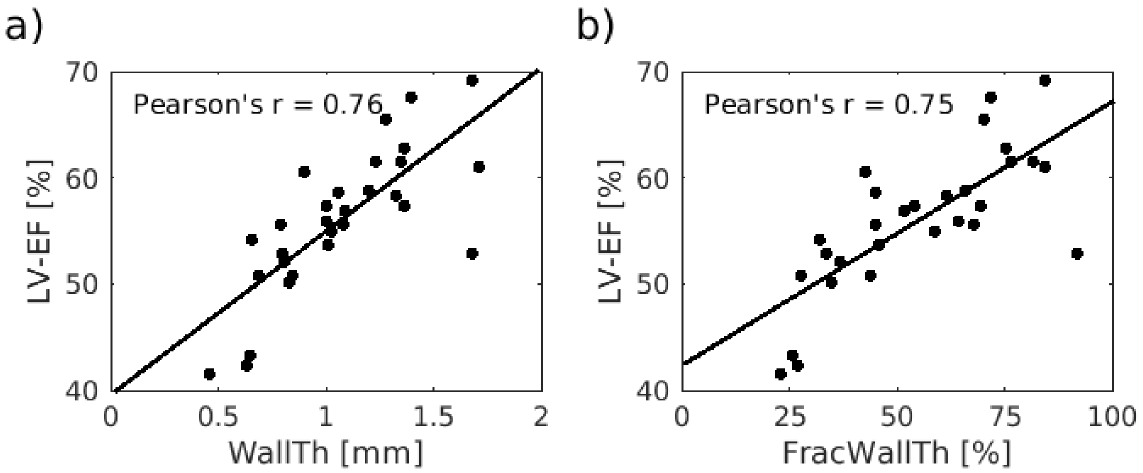


**Supplementary Figure 2: Myocardial segmentation.** A six sector segmentation model was used to calculate the left ventricular wall thickening of a mid-ventricular slice.

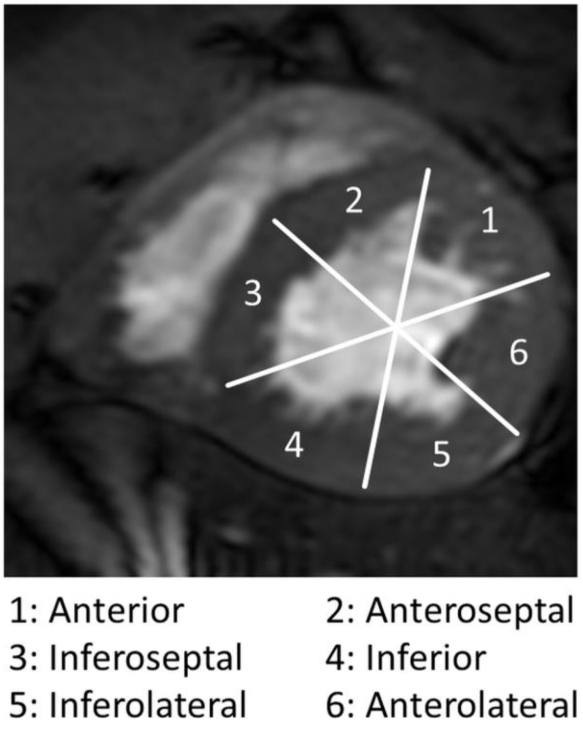

**Supplementary Table: Fractional wall thickening of the left ventricle itemized by segments**

| FracWallTh | Age groups | | | Female | Male (all) | All | p-value (one-way ANOVA, age) |
|---|---|---|---|---|---|---|---|
| | 23-58 Months | 63-106 Months | 110-168 Months | | | | |
| Anteroseptal | 41.0 ± 14.3 | 40.6 ± 15.6 | 36.2 ± 21.1 | 40.9 ± 19.6 | 37.7 ± 15.0 | 39.2 ± 17.4 | 0.82 |
| Inferoseptal | 57.4 ± 32.0 | 56.8 ± 25.5 | 60.3 ± 17.4 | 58.5 ± 30.4 | 57.9 ± 20.3 | 58.2 ± 25.7 | 0.96 |
| Inferior | 55.1 ± 27.4 | 52.3 ± 25.4 | 62.4 ± 18.1 | 58.3 ± 23.9 | 55.3 ± 24.5 | 56.7 ± 24.3 | 0.67 |
| Inferolateral | 73.9 ± 35.6 | 55.3 ± 30.3 | 59.9 ± 16.4 | 62.5 ± 28.0 | 64.0 ±31.0 | 63.3 ± 29.6 | 0.38 |
| Anterolateral | 76.0 ± 29.0 | 55.4 ± 33.1 | 57.9 ± 26.3 | 60.0 ± 29.0 | 66.5 ± 32.2 | 63.4 ± 30.9 | 0.29 |
| Anterior | 52.8 ± 16.3 | 46.6 ± 24.3 | 41.3 ± 18.1 | 46.5 ± 23.5 | 47.3 ± 16.7 | 46.9 ± 20.3 | 0.47 |

All data given as arithmetic mean and standard deviation. Significance level $p < 0.05$.
